# The in situ structures of mono-, di-, and tri-nucleosomes in human heterochromatin

**DOI:** 10.1101/334490

**Authors:** Shujun Cai, Désirée Böck, Martin Pilhofer, Lu Gan

**Author notes:** These authors contributed equally to this work (Shujun Cai, Désirée Böck).

## Abstract

The *in situ* 3-D organization of chromatin at the nucleosome and oligonucleosome levels is unknown. Here we use cryo-electron tomography (cryo-ET) to determine the *in situ* structures of HeLa nucleosomes, which have canonical core structures and asymmetric, flexible linker DNA. Subtomogram remapping suggests that sequential nucleosomes in heterochromatin follow irregular paths at the oligonucleosome level. This basic principle of higher-order repressive chromatin folding is compatible with the conformational variability of the two linker DNAs at the single-nucleosome level.

## INTRODUCTION

The fundamental unit of chromatin is the nucleosome, a 10-nm diameter, 6-nm thick cylindrical complex assembled from eight histone proteins and wrapped ∽1.65 times by 146 bp of DNA (Luger *et al.*, 1997; Chua *et al.*, 2016). In cells, many nucleosomes bind a linker histone, which stabilizes the two linker DNAs in a crossed conformation at the entry/exit position. When isolated or reconstituted, this larger nucleosome complex is called the chromatosome (Zhou *et al.*, 2015; Bednar *et al.*, 2017). Chemically fixed nucleosome chains can form highly ordered 30-nm fibers *in vitro* (Routh *et al.*, 2008; Song *et al.*, 2014), but these structures have not been detected inside cycling metazoan, plant, or yeast cells (McDowall *et al.*, 1986; Bouchet-Marquis *et al.*, 2006; Eltsov *et al.*, 2008; Fussner *et al.*, 2011; Fussner *et al.*, 2012; Gan *et al.*, 2013; Eltsov *et al.*, 2014; Chen *et al.*, 2016; Cai *et al.*, 2017; Ou *et al.*, 2017; Eltsov *et al.*, 2018). While the consensus is that *in situ* chromatin structure is irregular (Hansen *et al.*, 2018), the 3-D details of chromatin packing at the nucleosome level remain unknown.

Individual nucleosomes are challenging to identify *in situ*. The DNA-proximal negative-stain approach (ChromEMT) suffers distortions from chemical fixation, dehydration, and staining, resulting in large groups of nucleosomes appearing as amorphous elongated bodies instead of sets of discrete particles (Ou *et al.*, 2017). For cryo-EM samples, small high-contrast non-perturbative stains do not yet exist. Immuno-EM is also not suitable for protein identification because (1) antibodies would freeze immediately upon contact with any part of a cryo-EM sample, (2) the antibodies can only access the two surfaces of typical EM samples, and (3) antibody-gold complexes are too large (> 10 nm) to unambiguously identify small complexes in crowded environments. Correlative cryo-light/cryo-super-resolution microscopy can facilitate the localization of rare or sparsely distributed complexes, but it does not yet have sufficient resolution to identify individual nucleosomes in the crowded nucleoplasm (Chang *et al.*, 2014). Large multi-megadalton complexes have been successfully identified by their structural signature (their size and shape) (Medalia *et al.*, 2002). This non-invasive approach could in principle be done for smaller complexes like nucleosomes. We have therefore taken advantage of sample-preparation, imaging-hardware, and image-processing advances to determine the structures, positions, and orientations of nucleosomes inside a HeLa cell. Our resultant subtomogram averages and remapped models reveal a first glimpse of higher-order heterochromatin structure and folding up to the trinucleosome level *in situ*.

## RESULTS AND DISCUSSION

To determine how interphase mammalian chromatin is organized *in situ*, we performed Volta phase-contrast cryo-electron tomography (cryo-ET) (Fukuda *et al.*, 2015) on a HeLa cell that was thinned with a new cryo-focused-ion-beam (cryo-FIB) milling workflow (Medeiros *et al.*, 2018). The resultant cryotomogram shows exceptional detail, such as the clear delineation of membrane leaflets and a nucleoplasm densely populated with nucleosomes (Figure 1, A and B). Unlike interphase yeast cells, which have uniformly distributed nucleosomes (Chen *et al.*, 2016; Cai *et al.*, 2017), mammalian cells have densely packed perinuclear heterochromatin (Figure 1C) flanking the nuclear pore and loosely packed euchromatin (Figure 1D) in the interior positions (Visser *et al.*, 2000; van Steensel and Belmont, 2017).

**FIGURE 1:**
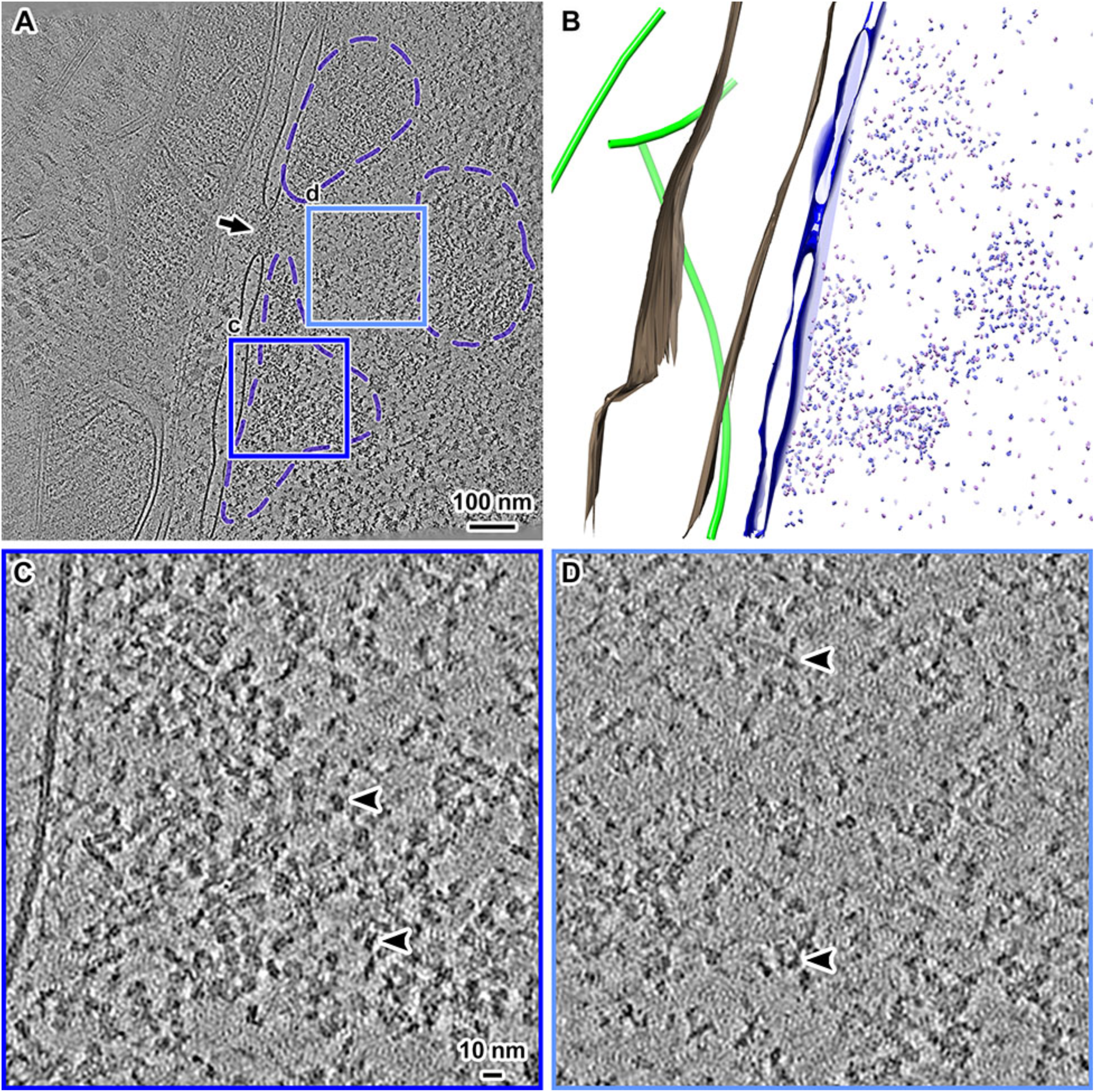
Volta cryotomogram of a cryo-FIB-thinned HeLa cell. **(A)** Tomographic slice (20 nm) of the nuclear periphery of a HeLa cell. The nuclear pore is indicated by the arrow. Three heterochromatic positions are delineated by purple dashed lines. (**B**) Segmentation of the mitochondrial outer membrane (brown), microtubules (green), and the nuclear envelope (dark blue). The *in silico* purified nucleosomes are also shown (blue and magenta puncta, see text). (**C**, **D**) Tomographic slices (10 nm) of the (**C**) heterochromatin and (**D**) euchromatin positions boxed in panel **A**, enlarged 4.5-fold. Several nucleosomes are indicated by arrowheads.

Multi-megadalton complexes are straightforward to identify in cryotomograms (Briegel *et al.*, 2009; Gan *et al.*, 2011; Xi *et al.*, 2011; Asano *et al.*, 2015; Mahamid *et al.*, 2016; Böck *et al.*, 2017), but nucleosomes are not because they are only ∽200 kilodaltons. In this study, we purify the nucleosomes “*in silico*” by combining template matching with 3-D classification (Bharat and Scheres, 2016), which we previously showed to be sensitive enough to identify nucleosomes of different linker-DNA conformations in nuclear lysates (Cai *et al.*, 2018). To minimize model bias, we used a featureless cylindrical reference (Figure 2A). This approach reveals a single 3-D class average that has the nucleosome’s unmistakable structural signatures: the left-handed wrapping of DNA, with the groove between the two gyres clearly visible at the position opposite of the DNA entry/exit site (Figure 2B). Like all crystal and cryo-EM structures, the nucleosome class average has two-fold symmetry around the dyad axis. Negative control 3-D classification of cytoplasmic densities that were template matched the same way did not produce any nucleosome-like class averages (Supplemental Figure S1).

**FIGURE 2:**
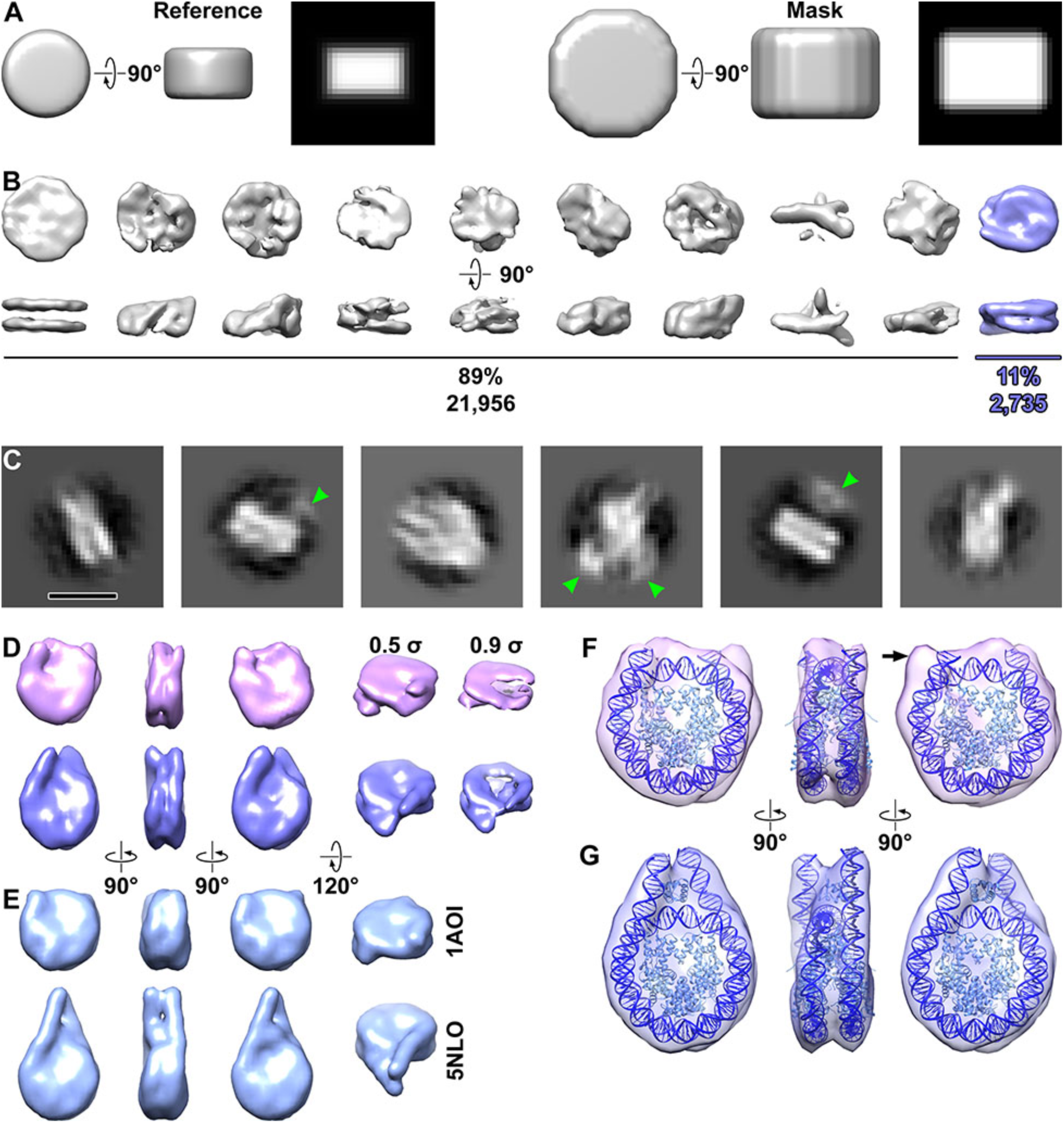
Structural analysis of nucleosomes *in situ*. (**A**) The reference model (left half) and mask (right half) used for 3-D classification and averaging. The reference is 10 nm wide and 6 nm thick. Both the reference and mask have soft edges that slowly decay to zero. The rightmost subpanel (black background) for both the reference and mask are central slices through the side view. (**B**) Three-dimensional class averages of all template-matching hits. The nucleosome class average (blue) is oriented with its two-fold dyad axis running horizontal. (**C**) Example 2-D class averages from the nucleosomes identified by 3-D classification. Some of the class averages that have densities from nucleosome-associated complexes (green arrowheads). Bar, 10 nm. (**D**) Final 3-D class averages of nucleosomes, showing from left to right the front, side, and back, and oblique views. One class (39%, magenta) has shorter linker-DNA densities that the other (61%, blue). Maps in all columns are contoured at 0.5 σ except in the rightmost column, which is set to 0.9 σ to better show the left-handed superhelical DNA path and the degree of linker DNA heterogeneity. (**E**) Crystal structures of the nucleosome core (PDB 1AOI, upper) and chromatosome (PDB 5NLO, lower), rendered as 15 Å resolution density maps. (**F,G**) Refined maps of the two nucleosome classes, with the edited chromatosome crystal structure docked (**F**) without or (**G**) with the linker histone, and linker DNA appropriately truncated. The histones and DNA are light and dark blue, respectively.

Unlike our previous analysis of picoplankton nuclear lysates in the which the nucleosomes were highly dispersed (Cai *et al.*, 2018), 3-D classification of HeLa nucleosomes in the crowded nucleus requires a cylindrical mask (Figure 2A). When we performed reference-free 2-D classification on the nucleosomes with a larger circular mask, we found that some class averages had extra densities in contact with the nucleosome (Figure 2C, green arrowheads). These densities are truncated by the mask, meaning that they belong to larger structures. Furthermore, these extra densities are weaker and featureless, consistent with their being averages of many different types of nucleosome-binding partners. Additional rounds of 3-D classification produced a final set of 1,141 nucleosomes (see Materials and Methods).

Three-dimensional classification of the final nucleosome set into two classes yielded averages with either short or long linker-DNA densities (Figure 2D). These two classes refined to 24 and 21 Å resolution, respectively (Supplemental Figure S2), and resemble low-pass-filtered density maps calculated from crystal structures (Figure 2E). Indeed, the averages can accommodate the chromatosome crystal structure after rigid-body alignment and adjustment of the linker DNA lengths (Figure 2, F and G) (Bednar *et al.*, 2017). The class with the shorter linker DNA can be best fit with a nucleosome core (151 bp). One of the linker-DNA densities cannot be adequately accounted for by the nucleosome crystal structure (Figure 2F and Supplemental Movie S1) and is instead consistent with partial unwrapping (Bilokapic *et al.*, 2018b, a). The nucleosome class with the longer linker DNA is best fit with ∽13 bp of DNA in each linker (172 bp total, Figure 2G and Supplemental Movie S2). The linker DNAs have a crossed conformation and remain visible when the density map’s contour level is raised (Figure 2D). This structural phenotype is consistent with the linker DNA’s conformational stabilization by a linker histone. Note that HeLa cells have a 183-bp nucleosome-repeat length (Lohr *et al.*, 1977), which predicts that sequential nucleosomes are linked by an average ∽12 nm DNA (37bp × 0.34 nm). Therefore, the short linker DNA densities in one of the subtomogram averages arises from linker-DNA conformational heterogeneity in the individual nucleosomes. Finally, classification of the 1,141 nucleosomes into four classes produces averages that show additional linker-DNA conformations, supporting the notion that the linker DNA is the most conformationally heterogeneous part of the nucleosome (Supplemental Figure S3).

Our subtomogram averages presented an opportunity to visualize nucleosomes in the context of higher-order chromatin structure *in situ*. Using the 3-D refined orientations and positions, we remapped the nucleosomes back into an empty volume the size of the original cryotomogram (Figure 3). As expected from their appearance in the tomographic slices (Figure 1A), the nucleosomes are predominantly localized in the three heterochromatin clusters (Figure 3A). The heterochromatin and euchromatin contain both classes of nucleosomes (Figure 3, C and D). The nucleosomes in between the heterochromatin domains appear isolated instead of being parts of contiguous chains. Some nucleosomes must have been missed by our analysis. For example, nucleosomes oriented with their face parallel to the lamella surface were missed (Supplemental Figure S2, B and C); nucleosomes oriented this way are known to be challenging to locate in plunge-frozen samples (Chua *et al.*, 2016). Our analysis would also have missed nucleosomes that make multiple contacts with large protein complexes (McGinty and Tan, 2015; Morgan *et al.*, 2016; Wilson *et al.*, 2016; Xu *et al.*, 2016; Farnung *et al.*, 2017; Liu *et al.*, 2017; Ayala *et al.*, 2018; Eustermann *et al.*, 2018) and nucleosomes with unconventional structures such as partially unwrapped nucleosomes (Bilokapic *et al.*, 2018a, b) and hexasomes (Kato *et al.*, 2017).

**FIGURE 3:**
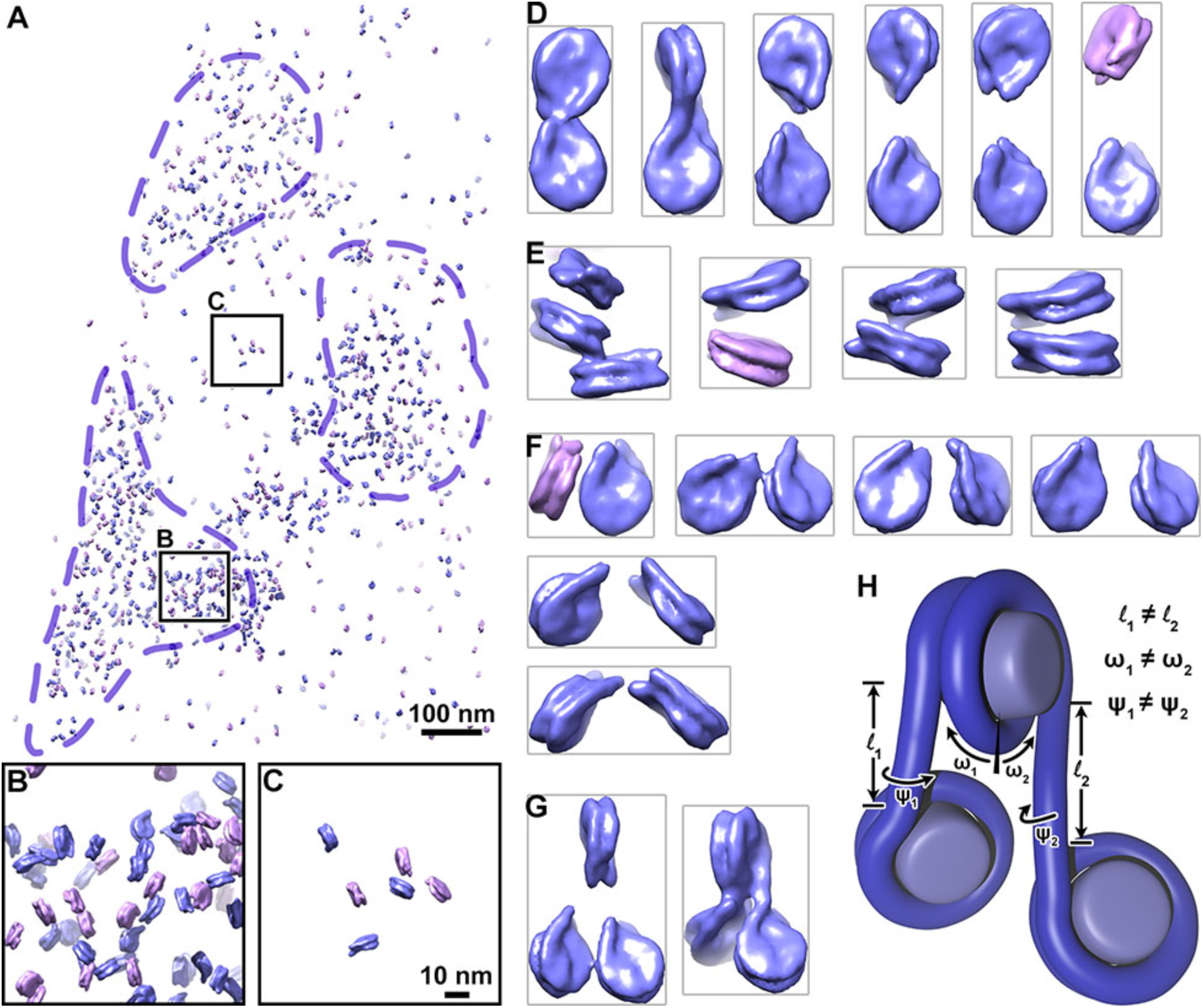
Chromatin is irregular at the oligonucleosome level *in situ*. (**A**) Model of short-linker (magenta) and long-linker (blue) nucleosomes remapped according to their positions and orientations in the nucleus. Dashed purple lines indicate approximate boundaries of heterochromatin. (**B, C**) Four-fold enlargements of the heterochromatin and a euchromatin positions boxed in panel **A**. (**D-G**) Examples of (**D**) dinucleosomes connected by linker DNA, (**E**) face-to-face packed nucleosomes, (**F**) dinucleosomes not connected by linker DNA but likely to be in sequence with a third nucleosome that was missed by our analysis, and (**G**) trinucleosomes connected by linker DNA. For clarity, adjacent remapped nucleosomes were cropped out. (**H**) The lengths (l_1_, l_2_), angles relative to the dyad axis (ω_1_, ω_2_), and rotation around the linker-DNA axes (ψ_1_, ψ_2_) are uncorrelated.

Many nucleosomes are likely to be interacting with each other because their linker-DNA densities are coaxial or because their cores are nearly stacked. We recognized four types of nucleosome-nucleosome arrangements (Figure 3, D-G and Supplemental Figure S4): nucleosome pairs likely to be connected by linker DNA (Figure 3D); nucleosome pairs oriented with face-to-face interactions (Figure 3E); nucleosome pairs likely to share linker DNA with a third, unmapped nucleosome (Figure 3F); and trinucleosomes likely connected by linker DNA (Figure 3G). The visualization of linker DNA densities in the subtomogram averages and remapped models provides the first clues about the path of DNA at the trinucleosome level (Figure 3, F and G). Nucleosomes in these examples are likely to follow an irregular zig-zag path. Periodic motifs such as those found in tetranucleosomes were not found and therefore must be exceptionally rare (Schalch *et al.*, 2005; Song *et al.*, 2014).

Chromatin higher-order structure is extremely sensitive to linker DNA parameters. For example, tetranucleosome face-to-face stacking can be abolished *in vitro* with a small change in linker-DNA length (Ekundayo *et al.*, 2017). Recent cryo-EM studies showed that dinucleosomes have variable conformations even when they are reconstituted with a strong positioning sequence and are bound to either heterochromatin protein 1 or Polycomb repressive complex 2 (Machida *et al.*, 2018; Poepsel *et al.*, 2018). Our subtomogram averages and remapped nucleosomes are consistent with a model in which variations of linker DNA length and orientation at the single nucleosome level *in situ* give rise to irregular higher-order chromatin structure at the dinucleosome and trinucleosome levels (Figure 3H). Chromatin can therefore pack densely in heterochromatin without folding into periodic motifs. Future advances in cryo thinning, automation, subtomogram classification, and remapping will be important tools to dissect *in situ* chromatin structure in greater detail.

## MATERIALS AND METHODS

### Cell culture

HeLa CCL2-cells (ATCC) were grown in DMEM (Gibco) supplemented with 10% inactivated FCS (Invitrogen) and 50 μg/mL streptomycin (AppliChem) at 37°C and 5% CO2. For electron microscopy (EM) imaging experiments, EM finder grids (gold NH2 R2/2, Quantifoil) were sterilized under UV light and then glow discharged. Grids were placed on the bottom of the wells of a 12-well plate (Nunc, Thermo Fisher) and equilibrated with DMEM for 30 min. Subsequently, 30,000 HeLa cells were seeded into each well and incubated overnight until vitrification.

### Preparation of frozen-hydrated specimens

Plunge freezing was performed as previously reported (Weiss *et al.*, 2017). Grids were removed from the wells using forceps. The forceps were then mounted in the Vitrobot and the grid was blotted from the backside by installing a Teflon sheet on one of the blotting pads. Grids were plunge-frozen in liquid ethane-propane (37 %/63 %) (Tivol *et al.*, 2008) using a Vitrobot Mk 4 (Thermo Fisher) and stored in liquid nitrogen.

### Cryo-FIB milling

Cryo-FIB was used to cryo-thin samples of plunge-frozen HeLa cells so that they could be imaged by cryo-ET (Marko *et al.*, 2007). Frozen grids with HeLa cells were first clipped into modified Autogrids (Thermo Fisher) (Medeiros *et al.*, 2018) and then transferred into the liquid-nitrogen bath of a loading station (Leica Microsystems). Grids were clamped onto a “40° pre-tilted TEM grid holder” (Leica Microsystems) and the holder was subsequently shuttled from the loading station to the dual-beam instrument using the VCT100 transfer system (Leica Microsystems). The holder was mounted on a custom-built cryo stage (Leica Microsystems) in a Helios NanoLab600i dual-beam FIB/SEM instrument (Thermo Fisher). The stage temperature was maintained below-154°C during the loading, milling and unloading procedures. Grid quality was checked by scanning EM imaging (5 kV, 21 pA). Samples were coated with a platinum precursor gas using the Gas Injector System and a “cold deposition” technique (Hayles *et al.*, 2007). Lamellae were milled in several steps. We first targeted two rectangular regions with the ion beam set to 30 kV and ∽400 pA to generate a ∽2-μm-thick lamella. The ion-beam current was then gradually decreased until the lamella reached a nominal thickness of ∽200 nm (ion beam set to ∽25 pA). After documentation of the lamellae by scanning EM imaging, the holder was brought back to the loading station using the VCT100 transfer system. The grids were unloaded and stored in liquid nitrogen.

### Electron cryomicroscopy and electron cryotomography

The cryo-EM imaging details are listed in Table S1. Cryo-FIB-processed HeLa cells were examined by both cryo-EM and cryo-ET (Weiss *et al.*, 2017). Images were recorded on a Titan Krios transmission electron cryomicroscope (Thermo Fisher) equipped with a K2 Summit direct-detection camera (Gatan), Quantum LS imaging filter (Gatan), and a Volta phase plate (Thermo Fisher). The microscope was operated at 300 kV with the imaging filter slit width set to 20 eV. Data were collected in-focus using the Volta phase plate. The pixel size at the specimen level was 3.45 Å. Tilt series covered an angular range from-60° to+60° with 2° increments. The total dose of a tilt series was 120 e^-^/Å^2^. Tilt series and 2-D projection images were acquired automatically using SerialEM(Mastronarde, 2005). Three-dimensional reconstructions and segmentations were generated using the IMOD program suite (Mastronarde, 2008). To increase the contrast, the tilt series was binned two-fold in the IMOD program *Etomo*, resulting in a final specimen-level pixel size of 6.9 Å.

### Template matching

The subtomogram analysis strategy was to find as many candidate nucleosomes as possible, then remove the majority of false positives by 3-D classification (Cai *et al.*, 2018). Template matching was done with PEET (Nicastro *et al.*, 2006; Heumann, 2016). To speed up the search, the tomogram was binned three-fold, corresponding to a 10.35 Å voxels. A featureless 10 nm diameter × 6 nm thick cylinder was created with the Bsoft (Heymann and Belnap, 2007) program *beditimg* and for use as the initial reference model. To emulate the effects of Volta phase contrast, this reference was corrupted with the Bsoft program *bctf* using a 3-D contrast transfer function with the fraction of amplitude contrast set to 0.5. To suppress the effects of nucleoplasmic background densities, the template was masked with a soft-edged cylinder. To minimize the number of false negatives, we used a very low cross-correlation cutoff of CC=0.2. We also set the minimum inter-particle spacing to 6 nm, which ensured that any face-to-face stacked nucleosomes would not be missed. To minimize model bias, only data up to ∽50 Å resolution were used. Using these criteria, ∽24,700 of ∽83,300 possible hits were retained. Visual inspection of the hits list confirmed that many non-nucleosome densities were also included.

### Classification analysis and 3-D subtomogram remapping

All 2-D and 3-D classification and 3-D auto-refinement were done with RELION 2.1 (Kimanius *et al.*, 2016) using default parameters except where noted below. A large box (∽2× the nucleosome diameter) was used so that the particle center could be refined during classification. This box choice resulted in the introduction of new false positives, which were dealt with a second round of template matching (see below). The template-matched particles were extracted using the subtomogram analysis routines (Bharat *et al.*, 2015). Orientation information was discarded in this process. For 2-D classification, the mask diameter was 140 Å and the regularization parameter T was set to 4. Three-dimensional classification was done with a featureless 10-nm diameter × 6-nm thick cylindrical reference and a larger cylindrical mask with a soft edge (Figure 2A).

Sequential rounds of 3-D classification pruned the nucleosome class to 1,883 particles. RELION performs classification and alignment simultaneously, meaning that it functions as another form of multi-class template matching in which the templates can change during the run. One consequence is that some nucleosome centers can translate to positions that either overlap neighboring nucleosomes or correspond to the nucleoplasm. To deal with the existence of new false positives, an additional round of template matching was performed in PEET, using only the refined positions of the 1,883 classified particles that contributed to the nucleosome class averages. The refined nucleosome density map (including the 1,883 particles) was used as the new template-matching reference. PEET removed the duplicated particles automatically. Next, the cross-correlation threshold relative to the template was incrementally increased until most of the spurious positions in the nucleoplasm were removed, yielding the final set of 1,141 nucleosomes. A final 3-D classification was performed with two classes, resulting in one class average with long linker DNA and one with short linker DNA. Following 3-D autorefinement, the angular distribution was checked by loading the final .bild file and density maps together in UCSF Chimera (Pettersen *et al.*, 2004).

The nucleosome averages were remapped using the script *ot_remap.py* (https://github.com/anaphaze/ot-tools), which orients and positions each RELION class average into an empty volume the same size as the original tomogram (Cai *et al.*, 2018). One remapped model was created for each class (short and long linker DNA). The two models were then combined with the Bsoft program *badd*. Because the pair-wise inter-nucleosome distances and positions, i.e., higher-order structure, was so heterogeneous, dinucleosomes and trinucleosomes had to be located manually in UCSF Chimera. To facilitate this manual search, the clipping planes were positioned so that the thickness along the view axis was < 40 nm. Pairs of nucleosomes were considered to be interacting if their linker DNAs were aligned (sequential nucleosomes) or if any part of the two nucleosomes were within ∽2 nm.

### Crystal structure docking

Because cryo-ET *in situ* subtomogram averages have much-lower resolutions than crystal structures, the goal was to conservatively dock a chromatosome crystal structure into the subtomogram averages. Of the two chromatosome structures (Zhou *et al.*, 2015; Bednar *et al.*, 2017), 5NL0 fit as a rigid body into the class with long linker DNA with minimal modification. This crystal structure was used as a starting point for further editing. For the nucleosome with longer linker DNA, 13 and 12 base pairs were removed from the linker-DNA termini, leaving 172 bp DNA. For the nucleosome with shorter linkers, 24 and 22 bp of DNA was removed from the linker-DNA termini, leaving 151 bp of DNA. Next the chromatosome model was docked automatically with the UCSF Chimera *fit-in-map* routine, using a map simulated to 20 Å resolution. These produced map-to-model correlations of 0.95 (nucleosome with long linker) and 0.87 (nucleosome with short linker). Owing to the limited resolution, no further attempts were made to refine the atomic model.

### Graphics

Figure panels were created in Adobe Illustrator CS6, Google Sheets, or Blender 2.79 (https://www.blender.org) and then arranged in Adobe Photoshop CS6.

## Data availability

The unbinned frame-aligned tilt series was deposited in the Electron Microscopy Public Image Archive (Iudin *et al.*, 2016) as EMPIAR-10179. The two-fold binned tomogram and the nucleosome subtomogram averages with short and long linker DNA were deposited in the Electron Microscopy Data Bank as EMD-6948, EMD-6949, and EMD-6950, respectively.

### ACKNOWLEDGEMENTS

We thank Duane Loh and Reza Khayat for discussions on heterogeneity and classification, John Heumann for advice on how to accelerate PEET template matching, and members of the Gan and Pilhofer teams and Alex Noble for feedback. ScopeM is acknowledged for instrument access at ETH Zürich. SC and LG were supported by a MOE T2 R-154-000-624-112, MOE T1 R-154-000-A49-114, and a NUS YIA R-154-000-558-133. DB and MP were supported by the European Research Council, the Swiss National Science Foundation, and the Helmut Horten Foundation.

## Contributions

SC-experiments, writing, DB-experiments, writing, MP-writing, LG-experiments, writing.

---

**SUPPLEMENTAL FIGURE S1:**
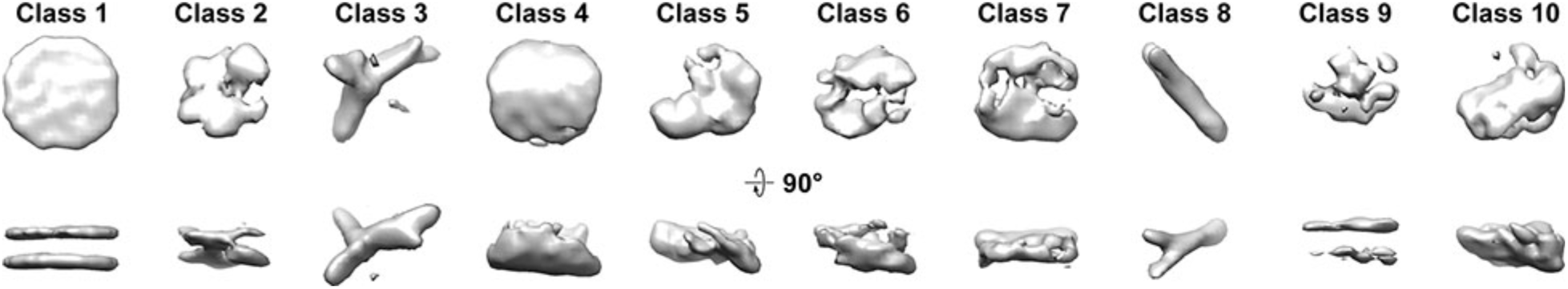
Classification control. Control 3-D class averages of “nucleosome” template-matching hits taken from the cytoplasm and mitochondrion.

**SUPPLEMENTAL FIGURE S2:**
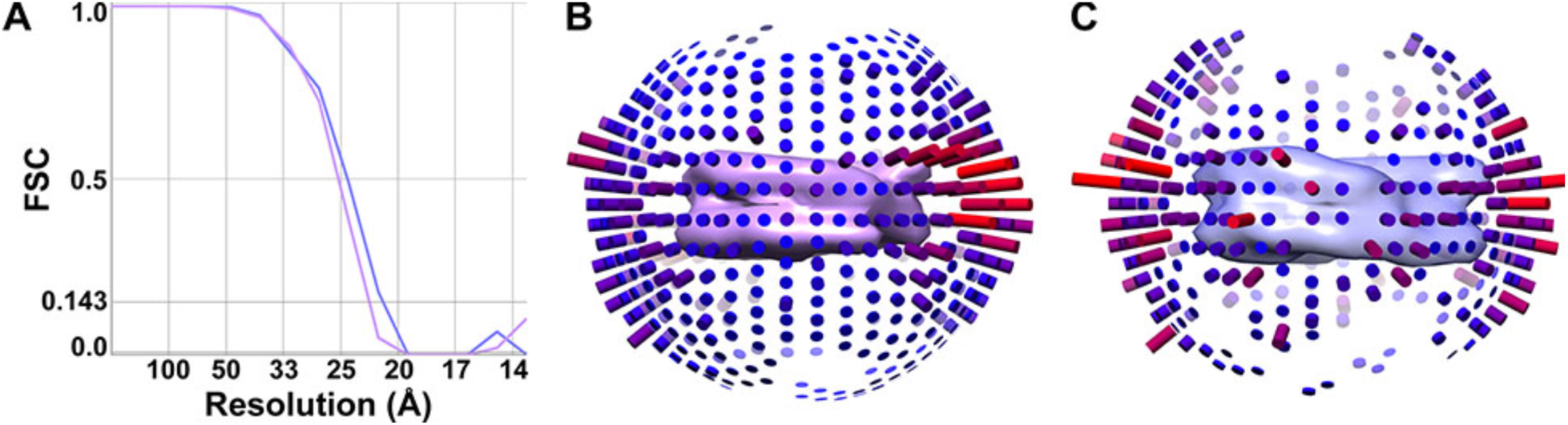
Analysis of nucleosome averages. (**A**), The resolution of the two nucleosome classes with short (magenta) and long (blue) linker DNA are respectively 24 and 21 Å based on the Fourier Shell Correlation (FSC)=0.143 criterion. (**B, C**) Angular-distribution plots of the nucleosome classes with (**B**) short linker DNA and (**C**) long linker DNA densities. The number of particles oriented with the view vector parallel to each cylinder is proportional to the cylinder’s height and redness. An isosurface of each average is shown at the center of the corresponding angular-distribution plot.

**SUPPLEMENTAL FIGURE S3:**
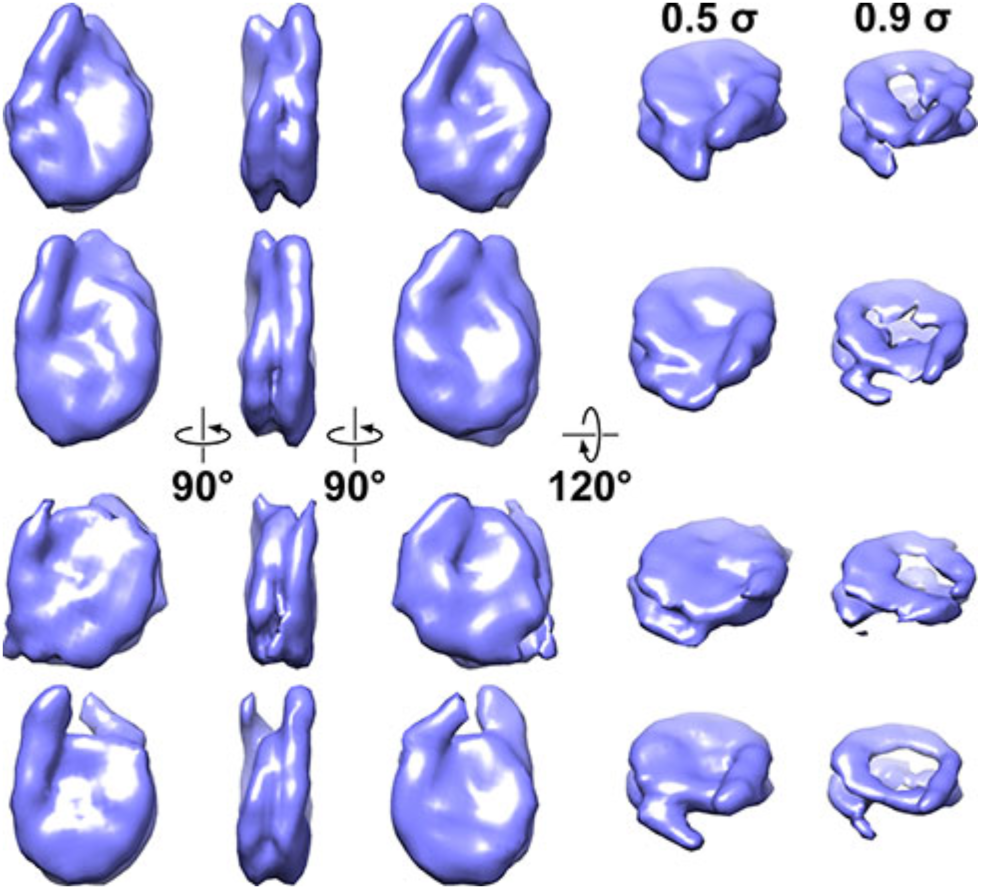
Three-dimensional classification into four classes. Classification of nucleosomes into four classes, showing the front, side, and back, and oblique views (left to right). All columns are presented as in Figure 2D.

**SUPPLEMENTAL FIGURE S4:**
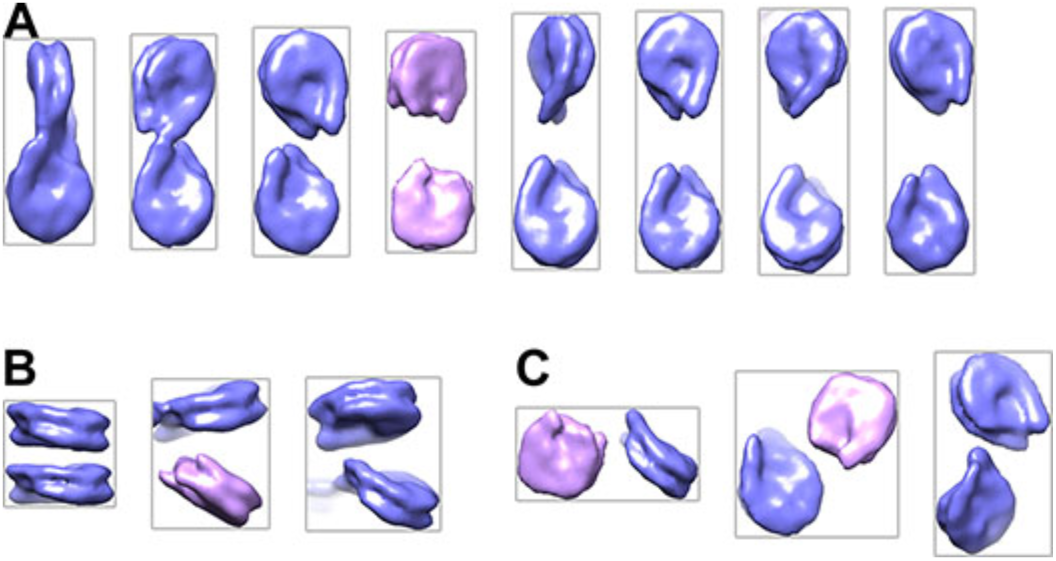
Additional examples of dinucleosomes. (**A**) Dinucleosomes that are likely to be connected by linker DNA. (**B**) Dinucleosomes interacting face to face or with their dyad axes intersecting at the left. (**C**) Dinucleosomes that are neither connected by linker DNA nor packed face to face. All panels share the color scheme as Figure 3.

**Supplemental Table S1:** **Cryo-ET parameters**

|  |  |
| --- | --- |
| Microscope | Titan Krios (Thermo Fisher) |
| Energy | 300 KeV |
| Gun type | FEG |
| Camera | K2 Summit (Gatan) |
| Recording mode | counting mode |
| Subframe duration | 0.4 seconds |
| Subframes / tilt | 4 |
| Energy filter | Quantum LS (Gatan) |
| Zero-loss slit width | 20 eV |
| Calibrated pixel size | 3.45 Å |
| Defocus | in focus, Volta phase contrast |
| Cumulative dose | 120 e<sup>-</sup> / Å<sup>2</sup> |
| Tilt range | ± 60°, bidirectional |
| Tilt increment | 2° |
| Tomography software | SerialEM 3.6.4 |
| Subframe alignment software | alignframes (IMOD) |
| Tomogram processing & visualization | IMOD 4.9 |
| Fiducial model generation | patch tracking (IMOD) |
| Template matching | PEET 1.11 |
| Density map creation & editing | Bsoft 1.8.8 |
| Subtomogram classification & averaging | RELION 2.1 |
| Density map visualization & docking | UCSF Chimera 1.11 |

## SUPPLEMENTAL MOVIES S1 and S2

**Density maps and docked atomic models of HeLa nucleosomes**

Subtomogram averages and models of the nucleosome classes with short (S1) and long (S2) linker DNAs are contoured at 0.5 σ. The color scheme is identical to Figure 2, F and G.

## REFERENCES

Asano, S., Fukuda, Y., Beck, F., Aufderheide, A., Forster, F., Danev, R., and Baumeister, W. (2015). Proteasomes. A molecular census of 26S proteasomes in intact neurons. Science 347, 439–442.

Ayala, R., Willhoft, O., Aramayo, R.J., Wilkinson, M., McCormack, E.A., Ocloo, L., Wigley, D.B., and Zhang, X. (2018). Structure and regulation of the human INO80-nucleosome complex. Nature.

Bednar, J., Garcia-Saez, I., Boopathi, R., Cutter, A.R., Papai, G., Reymer, A., Syed, S.H., Lone, I.N., Tonchev, O., Crucifix, C., Menoni, H., Papin, C., Skoufias, D.A., Kurumizaka, H., Lavery, R., Hamiche, A., Hayes, J.J., Schultz, P., Angelov, D., Petosa, C., and Dimitrov, S. (2017). Structure and Dynamics of a 197 bp Nucleosome in Complex with Linker Histone H1. Mol Cell 66, 384–397 e388.

Bharat, T.A., Russo, C.J., Lowe, J., Passmore, L.A., and Scheres, S.H. (2015). Advances in Single-Particle Electron Cryomicroscopy Structure Determination applied to Sub-tomogram Averaging. Structure 23, 1743–1753.

Bharat, T.A., and Scheres, S.H. (2016). Resolving macromolecular structures from electron cryo-tomography data using subtomogram averaging in RELION. Nat Protoc 11, 2054–2065.

Bilokapic, S., Strauss, M., and Halic, M. (2018a). Histone octamer rearranges to adapt to DNA unwrapping. Nat Struct Mol Biol 25, 101–108.

Bilokapic, S., Strauss, M., and Halic, M. (2018b). Structural rearrangements of the histone octamer translocate DNA. Nat Commun 9, 1330.

Böck, D., Medeiros, J.M., Tsao, H.F., Penz, T., Weiss, G.L., Aistleitner, K., Horn, M., and Pilhofer, M. (2017). In situ architecture, function, and evolution of a contractile injection system. Science 357, 713–717.

Bouchet-Marquis, C., Dubochet, J., and Fakan, S. (2006). Cryoelectron microscopy of vitrified sections: a new challenge for the analysis of functional nuclear architecture. Histochem Cell Biol 125, 43–51.

Briegel, A., Ortega, D.R., Tocheva, E.I., Wuichet, K., Li, Z., Chen, S., Muller, A., Iancu, C.V., Murphy, G.E., Dobro, M.J., Zhulin, I.B., and Jensen, G.J. (2009). Universal architecture of bacterial chemoreceptor arrays. Proc Natl Acad Sci U S A 106, 17181–17186.

Cai, S., Chen, C., Tan, Z.Y., Huang, Y., Shi, J., and Gan, L. (2017). Cryo-ET reveals nucleosome reorganization in condensed mitotic chromosomes in vivo. bioRxiv.

Cai, S., Song, Y., Chen, C., Shi, J., and Gan, L. (2018). Natural chromatin is heterogeneous and self-associates in vitro. Mol Biol Cell, mbcE17070449.

Chang, Y.W., Chen, S., Tocheva, E.I., Treuner-Lange, A., Lobach, S., Sogaard-Andersen, L., and Jensen, G.J. (2014). Correlated cryogenic photoactivated localization microscopy and cryo-electron tomography. Nat Methods 11, 737–739.

Chen, C., Lim, H.H., Shi, J., Tamura, S., Maeshima, K., Surana, U., and Gan, L. (2016). Budding yeast chromatin is dispersed in a crowded nucleoplasm in vivo. Mol Biol Cell 27, 3357– 3368.

Chua, E.Y., Vogirala, V.K., Inian, O., Wong, A.S., Nordenskiold, L., Plitzko, J.M., Danev, R., and Sandin, S. (2016). 3.9 A structure of the nucleosome core particle determined by phase-plate cryo-EM. Nucleic Acids Res 44, 8013–8019.

Ekundayo, B., Richmond, T.J., and Schalch, T. (2017). Capturing Structural Heterogeneity in Chromatin Fibers. J Mol Biol 429, 3031–3042.

Eltsov, M., Grewe, D., Lemercier, N., Frangakis, A., Livolant, F., and Leforestier, A. (2018). Nucleosome conformational variability in solution and in interphase nuclei evidenced by cryo-electron miocroscopy of vitreous sections. bioRxiv.

Eltsov, M., Maclellan, K.M., Maeshima, K., Frangakis, A.S., and Dubochet, J. (2008). Analysis of cryo-electron microscopy images does not support the existence of 30-nm chromatin fibers in mitotic chromosomes in situ. Proc Natl Acad Sci U S A 105, 19732–19737.

Eltsov, M., Sosnovski, S., Olins, A.L., and Olins, D.E. (2014). ELCS in ice: cryo-electron microscopy of nuclear envelope-limited chromatin sheets. Chromosoma 123, 303–312.

Eustermann, S., Schall, K., Kostrewa, D., Lakomek, K., Strauss, M., Moldt, M., and Hopfner, K.P. (2018). Structural basis for ATP-dependent chromatin remodelling by the INO80 complex. Nature.

Farnung, L., Vos, S.M., Wigge, C., and Cramer, P. (2017). Nucleosome-Chd1 structure and implications for chromatin remodelling. Nature 550, 539–542.

Fukuda, Y., Laugks, U., Lucic, V., Baumeister, W., and Danev, R. (2015). Electron cryotomography of vitrified cells with a Volta phase plate. J Struct Biol 190, 143–154.

Fussner, E., Djuric, U., Strauss, M., Hotta, A., Perez-Iratxeta, C., Lanner, F., Dilworth, F.J., Ellis, J., and Bazett-Jones, D.P. (2011). Constitutive heterochromatin reorganization during somatic cell reprogramming. EMBO J 30, 1778–1789.

Fussner, E., Strauss, M., Djuric, U., Li, R., Ahmed, K., Hart, M., Ellis, J., and Bazett-Jones, D.P. (2012). Open and closed domains in the mouse genome are configured as 10-nm chromatin fibres. EMBO Rep 13, 992–996.

Gan, L., Ladinsky, M.S., and Jensen, G.J. (2011). Organization of the smallest eukaryotic spindle. Curr Biol 21, 1578–1583.

Gan, L., Ladinsky, M.S., and Jensen, G.J. (2013). Chromatin in a marine picoeukaryote is a disordered assemblage of nucleosomes. Chromosoma 122, 377–386.

Hansen, J.C., Connolly, M., McDonald, C.J., Pan, A., Pryamkova, A., Ray, K., Seidel, E., Tamura, S., Rogge, R., and Maeshima, K. (2018). The 10-nm chromatin fiber and its relationship to interphase chromosome organization. Biochem Soc Trans 46, 67–76.

Hayles, M.F., Stokes, D.J., Phifer, D., and Findlay, K.C. (2007). A technique for improved focused ion beam milling of cryo-prepared life science specimens. J Microsc 226, 263–269.

Heumann, J.M. (2016). PEET: University of Colorado Boulder.

Heymann, J.B., and Belnap, D.M. (2007). Bsoft: image processing and molecular modeling for electron microscopy. J Struct Biol 157, 3–18.

Iudin, A., Korir, P.K., Salavert-Torres, J., Kleywegt, G.J., and Patwardhan, A. (2016). EMPIAR: a public archive for raw electron microscopy image data. Nat Methods 13, 387–388.

Kato, D., Osakabe, A., Arimura, Y., Mizukami, Y., Horikoshi, N., Saikusa, K., Akashi, S., Nishimura, Y., Park, S.Y., Nogami, J., Maehara, K., Ohkawa, Y., Matsumoto, A., Kono, H., Inoue, R., Sugiyama, M., and Kurumizaka, H. (2017). Crystal structure of the overlapping dinucleosome composed of hexasome and octasome. Science 356, 205–208.

Kimanius, D., Forsberg, B.O., Scheres, S.H., and Lindahl, E. (2016). Accelerated cryo-EM structure determination with parallelisation using GPUs in RELION-2. Elife 5.

Liu, X., Li, M., Xia, X., Li, X., and Chen, Z. (2017). Mechanism of chromatin remodelling revealed by the Snf2-nucleosome structure. Nature 544, 440–445.

Lohr, D., Corden, J., Tatchell, K., Kovacic, R.T., and Van Holde, K.E. (1977). Comparative subunit structure of HeLa, yeast, and chicken erythrocyte chromatin. Proc Natl Acad Sci U S A 74, 79–83.

Luger, K., Mader, A.W., Richmond, R.K., Sargent, D.F., and Richmond, T.J. (1997). Crystal structure of the nucleosome core particle at 2.8 A resolution. Nature 389, 251–260.

Machida, S., Takizawa, Y., Ishimaru, M., Sugita, Y., Sekine, S., Nakayama, J.I., Wolf, M., and Kurumizaka, H. (2018). Structural Basis of Heterochromatin Formation by Human HP1. Mol Cell.

Mahamid, J., Pfeffer, S., Schaffer, M., Villa, E., Danev, R., Cuellar, L.K., Forster, F., Hyman, A.A., Plitzko, J.M., and Baumeister, W. (2016). Visualizing the molecular sociology at the HeLa cell nuclear periphery. Science 351, 969–972.

Marko, M., Hsieh, C., Schalek, R., Frank, J., and Mannella, C. (2007). Focused-ion-beam thinning of frozen-hydrated biological specimens for cryo-electron microscopy. Nat Methods 4, 215–217.

Mastronarde, D.N. (2005). Automated electron microscope tomography using robust prediction of specimen movements. J Struct Biol 152, 36–51.

Mastronarde, D.N. (2008). Correction for non-perpendicularity of beam and tilt axis in tomographic reconstructions with the IMOD package. J Microsc 230, 212–217.

McDowall, A.W., Smith, J.M., and Dubochet, J. (1986). Cryo-electron microscopy of vitrified chromosomes in situ. EMBO J 5, 1395–1402.

McGinty, R.K., and Tan, S. (2015). Nucleosome structure and function. Chem Rev 115, 2255– 2273.

Medalia, O., Weber, I., Frangakis, A.S., Nicastro, D., Gerisch, G., and Baumeister, W. (2002). Macromolecular architecture in eukaryotic cells visualized by cryoelectron tomography. Science 298, 1209–1213.

Medeiros, J.M., Bock, D., Weiss, G.L., Kooger, R., Wepf, R.A., and Pilhofer, M. (2018). Robust workflow and instrumentation for cryo-focused ion beam milling of samples for electron cryotomography. Ultramicroscopy 190, 1–11.

Morgan, M.T., Haj-Yahya, M., Ringel, A.E., Bandi, P., Brik, A., and Wolberger, C. (2016). Structural basis for histone H2B deubiquitination by the SAGA DUB module. Science 351, 725– 728.

Nicastro, D., Schwartz, C., Pierson, J., Gaudette, R., Porter, M.E., and McIntosh, J.R. (2006). The molecular architecture of axonemes revealed by cryoelectron tomography. Science 313, 944–948.

Ou, H.D., Phan, S., Deerinck, T.J., Thor, A., Ellisman, M.H., and O’Shea, C.C. (2017). ChromEMT: Visualizing 3D chromatin structure and compaction in interphase and mitotic cells. Science 357.

Pettersen, E.F., Goddard, T.D., Huang, C.C., Couch, G.S., Greenblatt, D.M., Meng, E.C., and Ferrin, T.E. (2004). UCSF Chimera--a visualization system for exploratory research and analysis. J Comput Chem 25, 1605–1612.

Poepsel, S., Kasinath, V., and Nogales, E. (2018). Cryo-EM structures of PRC2 simultaneously engaged with two functionally distinct nucleosomes. Nat Struct Mol Biol 25, 154–162.

Routh, A., Sandin, S., and Rhodes, D. (2008). Nucleosome repeat length and linker histone stoichiometry determine chromatin fiber structure. Proc Natl Acad Sci U S A 105, 8872–8877.

Schalch, T., Duda, S., Sargent, D.F., and Richmond, T.J. (2005). X-ray structure of a tetranucleosome and its implications for the chromatin fibre. Nature 436, 138–141.

Song, F., Chen, P., Sun, D., Wang, M., Dong, L., Liang, D., Xu, R.M., Zhu, P., and Li, G. (2014). Cryo-EM study of the chromatin fiber reveals a double helix twisted by tetranucleosomal units. Science 344, 376–380.

Tivol, W.F., Briegel, A., and Jensen, G.J. (2008). An improved cryogen for plunge freezing. Microsc Microanal 14, 375–379.

van Steensel, B., and Belmont, A.S. (2017). Lamina-Associated Domains: Links with Chromosome Architecture, Heterochromatin, and Gene Repression. Cell 169, 780–791.

Visser, A.E., Jaunin, F., Fakan, S., and Aten, J.A. (2000). High resolution analysis of interphase chromosome domains. J Cell Sci 113 (Pt 14), 2585–2593.

Weiss, G.L., Medeiros, J.M., and Pilhofer, M. (2017). In Situ Imaging of Bacterial Secretion Systems by Electron Cryotomography. Methods Mol Biol 1615, 353–375.

Wilson, M.D., Benlekbir, S., Fradet-Turcotte, A., Sherker, A., Julien, J.P., McEwan, A., Noordermeer, S.M., Sicheri, F., Rubinstein, J.L., and Durocher, D. (2016). The structural basis of modified nucleosome recognition by 53BP1. Nature 536, 100–103.

Xi, Y., Yao, J., Chen, R., Li, W., and He, X. (2011). Nucleosome fragility reveals novel functional states of chromatin and poises genes for activation. Genome Res 21, 718–724.

Xu, P., Li, C., Chen, Z., Jiang, S., Fan, S., Wang, J., Dai, J., Zhu, P., and Chen, Z. (2016). The NuA4 Core Complex Acetylates Nucleosomal Histone H4 through a Double Recognition Mechanism. Mol Cell 63, 965–975.

Zhou, B.R., Jiang, J., Feng, H., Ghirlando, R., Xiao, T.S., and Bai, Y. (2015). Structural Mechanisms of Nucleosome Recognition by Linker Histones. Mol Cell 59, 628–638.

